# Surveying non-visual arrestins reveals allosteric interactions between functional sites

**DOI:** 10.1101/2022.05.20.492847

**Authors:** James M. Seckler, Emily N. Robinson, Stephen J. Lewis, Alan Grossfield

**Affiliations:** Department of Biomedical Engineering, Case Western Reserve University, Cleveland, Ohio, USA; Department of Biochemistry and Biophysics, University of Rochester, Rochester, NY, USA; Department of Pediatrics, Case Western Reserve University, Cleveland, Ohio, USA

**Keywords:** Anisotropic network modelling, arrestin, allostery, GPCR

## Abstract

Arrestins are important scaffolding proteins that are expressed in all vertebrate animals. They regulate cell signaling events upon binding to active G-protein coupled receptors (**GPCR**) and trigger endocytosis of active GPCRs. While many of the functional sites on arrestins have been characterized, the question of how these sites interact is unanswered. We used anisotropic network modelling (**ANM**) together with our covariance compliment techniques to survey all of the available structures of the non-visual arrestins to map how structural changes and protein-binding affect their structural dynamics. We found that activation and clathrin binding have a marked effect on arrestin dynamics, and that these dynamics changes are localized to a small number of distant functional sites. These sites include α-helix 1, the lariat loop, nuclear localization domain, and the C-domain β-sheets on the C-loop side. Our techniques suggest that clathrin binding and/or GPCR activation of arrestin perturb the dynamics of these sites independent of structural changes.

## 2. Introduction

The arrestin family of proteins has four members: the visual arrestins arrestin-1 and arrestin-4, and arrestin-2 and arrestin-3, also called β-arrestins 1 and 2 or non-visual arrestins [1, 2]. The first role identified for arrestins was the regulation of G-protein coupled receptor (**GPCR**) activity by triggering receptor endocytosis and recycling.[1, 2] More recent work has shown that non-visual arrestins also activate cell signaling proteins, [1, 2] including kinases such as mitogen-activated protein kinase (**MAPK**) and c-Jun N-terminal kinases (**JNK**). As such, arrestins are a second branch of GPCR signaling pathways that complement the primary G-protein pathway [3–7].

Arrestins are recruited to activated GPCRs after G-protein coupled receptor kinases (**GRK**) have phosphorylated the C-terminal tail of the GPCR [8], which triggers arrestin binding and activation. Upon activation, arrestin recruits clathrin and adaptin, triggering endocytosis of both the GPCR and arrestin [9–12]. Arrestin will then sort the bound GPCR into either a degradation or recycling pathway [13]. Afterwards, arrestin is translocated to the nucleus where it alters gene transcription [14]. The identity of the GPCR and the pattern of phosphorylation determines which of its downstream partners arrestin recruits [8, 15]. These factors control whether the GPCR is recycled or degraded, the specific kinases arrestin recruits, and which downstream effectors are triggered [8, 16–20]. The mechanism by which this process occurs remain elusive. We *hypothesize* that allosteric coupling of specific functional sites on arrestin convey the information about arrestin’s current binding partners and hence its final destiny in the cell.

All arrestin proteins consist of three distinct structural domains (**Figure 1**) [13]: the N-domain, the C-domain, and the C-tail. The N-domain is the N-terminal portion of the protein and consists of the GPCR binding site, α-helix 1, and residues D26 and R169 of the polar core [13, 21–23]. The C-domain forms the other half of the structured part of arrestin; this domain contains a pair of β-sheets, the C-loops, the Lariat loop, and residues D290 and D297 of the polar core [23]. The C-tail is a long unstructured loop that binds to the N-domain when arrestin is inactive; it contains residue R393 of the polar core and the clathrin binding site [13, 23]. The polar core of arrestin (D26, R169, D290, D297, and R393 in arrestin-2) is responsible for keeping it inactive when not bound to a GPCR. When arrestin binds a GPCR, the GPCR’s C-terminal tail displaces arrestin’s C-tail domain and the phospho-residues disrupt the charge balance of the polar core, allowing for activation (**Figure 2**)[24–26]. Upon arrestin activation, α-helix 1 stabilizes GPCR binding to arrestin without changing its position with respect to the rest of the N-domain [22], and the C-loops in the C-domain stabilize membrane binding upon arrestin activation [9, 10, 27]; this region is also responsible for binding clathrin. This leads to an interesting question: how do α-helix 1 and the C-loops only stabilize binding of their respective partners when arrestin is in the active form without changing their structure relative to the rest of the protein? Despite the wealth of crystallographic data, many questions remain about arrestin structure/function relationships.

**Figure 1:**
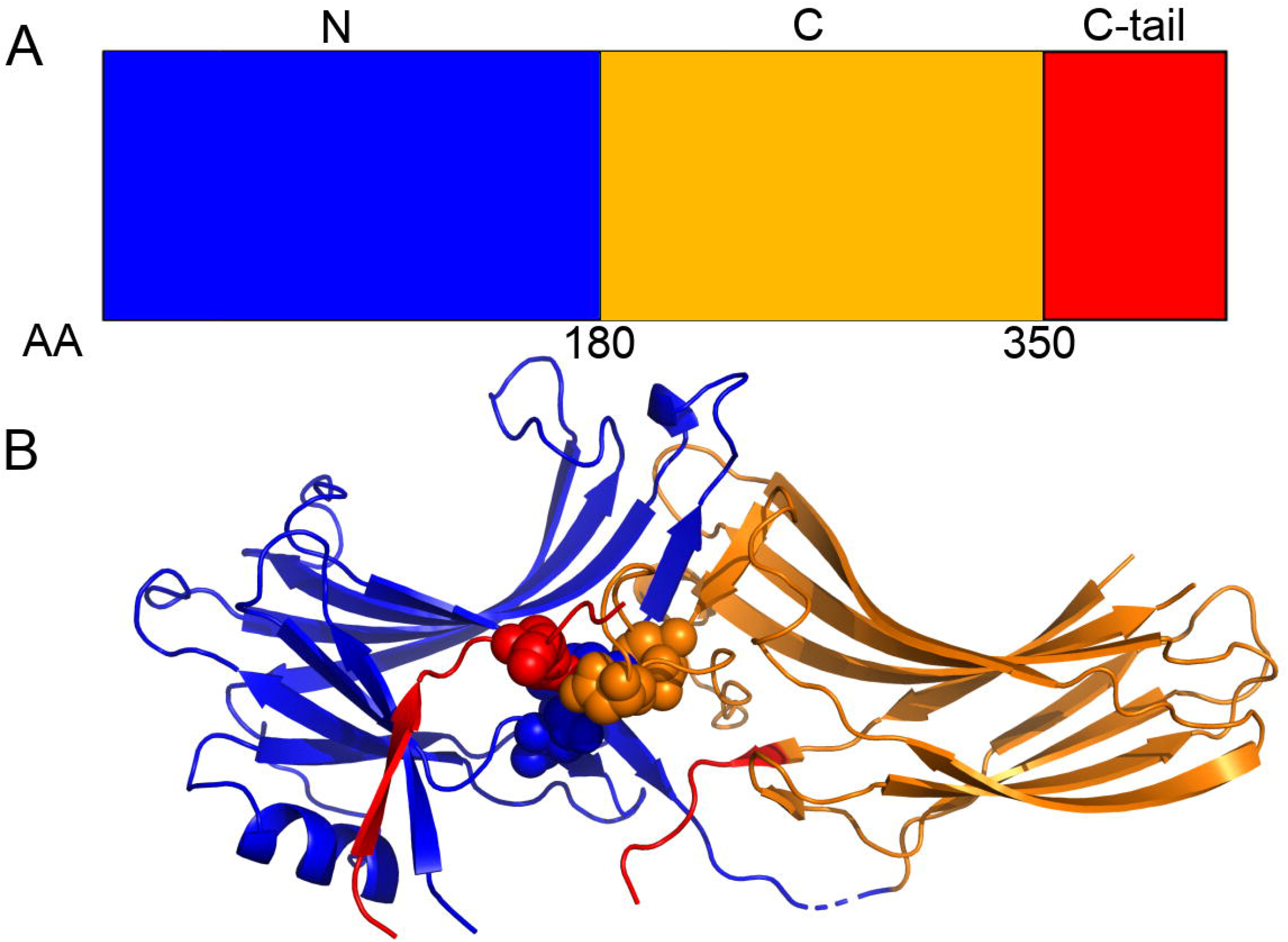
The A) domain and B) crystallography structure of Arrestin-2. The N-domain (blue), C-domain (orange), and C-tail domain (red) are shown along with the polar core domain (spheres). The discontinuity in the red backbone exists because a ~35-residue region of the C-tail domain is missing from all crystal structures.

**Figure 2:**
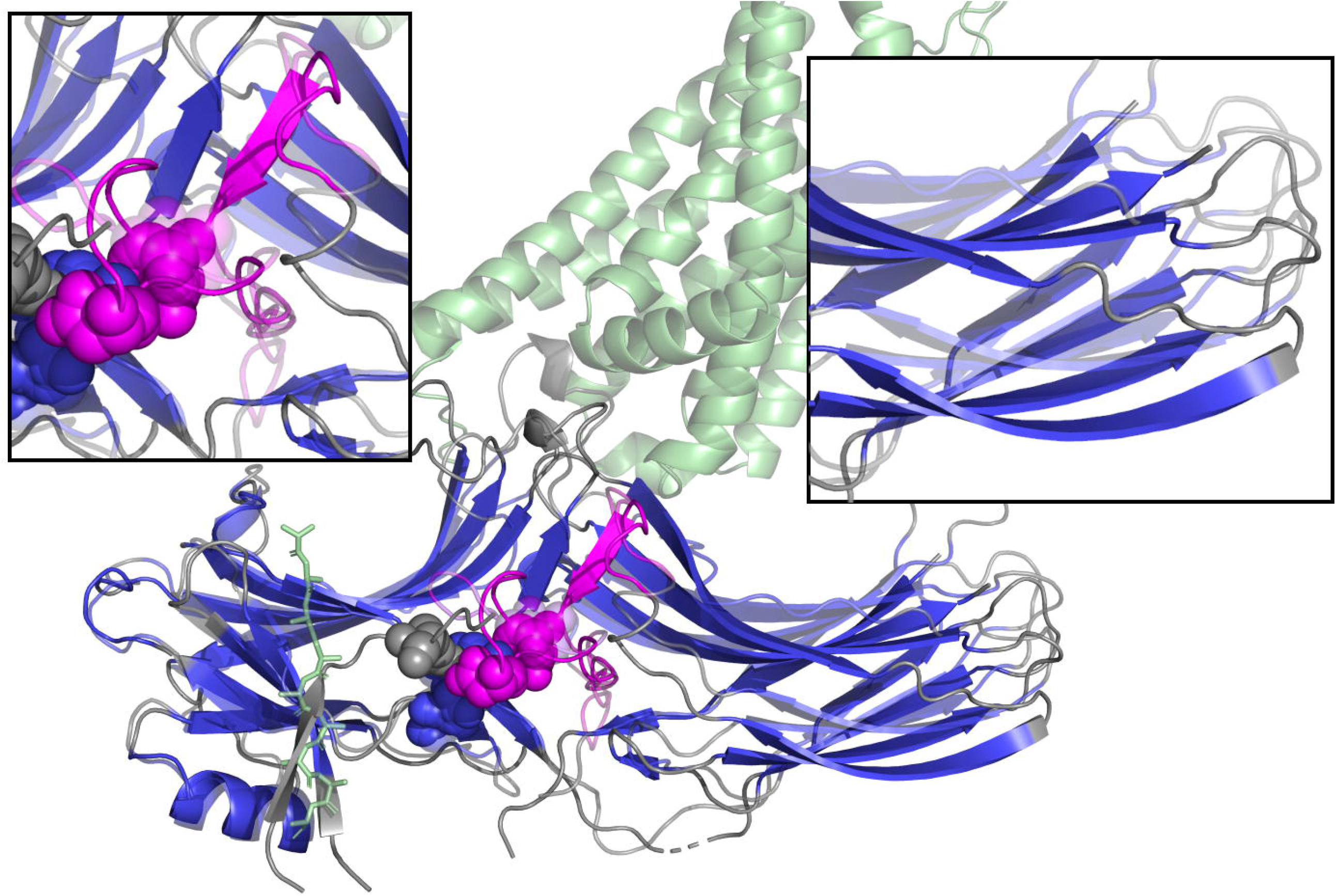
Changes in the structure of arrestin upon activation. Inactive arrestin (solid) and active arrestin (translucent) are superimposed on touch of each other, with the N-terminal domain on the left as in Figure 1. The polar core (spheres), lariat loop (purple), GPCR (green), and GPCR C-terminal peptide (green sticks) are shown. Blue and purple regions represent sequence which is included in our ANM modelling. Arrestin activation is marked by the disruption of the polar core residues (top left panel), and a rotation of the C-domain with respect to the N-domain (top right panel).

Elastic network models, specifically the anisotropic network model (**ANM**), are a valuable tool for rapidly probing large-scale motions of proteins[28–31]. In contrast to all-atom molecular dynamics simulations, which require a complex force field with many parameters and computationally expensive sampling[32], ANM uses a simple harmonic model of protein motions, where neighboring residues interact using simple springs. The fluctuations of the system are then obtained using an eigenvalue decomposition[28]. The resulting calculation is computationally efficient enough to be applied in an informatics setting, surveying all structures in a family [31]. Despite its simplicity, predictions from the low-frequency (high-amplitude) modes agree well with the dominant modes of principal component analysis of all-atom molecular dynamics simulations and with hydrogen deuterium exchange mass spectrometry (**HX-MS**), a technique for probing structural dynamics of *in vitro* proteins [30, 33–35].

Here we report that α-helix 1, the C-domain β-sheets on the C-loop side, the center of the lariat loop, and the nuclear localization sequence (**NLS**) alter their dynamic motions based on arrestin’s current state and binding partners. This provides a mechanism for how arrestin’s function changes during its activation and sorting process.

## 3. Methods

### 3.1 X-ray Structure Selection and Analysis

Crystallographic data were obtained from the Protein Data Bank [36, 37]. All apo, peptide-bound, and GPCR-bound X-ray structures of arrestin-2 and arrestin-3 were initially selected. If multiple protein chains were resolved in a single asymmetric unit, we separated these structures and analyzed them separately; these structures are denoted by their PDB id code with a (b) afterwards. We aligned the sequences and structures of these proteins and identified regions that were not resolved in the structures. We excluded two structures (3GC3, 6KL7(b)) because too many residues were not resolved. We identified the regions that were present in all structures and focused our analysis on that core. In addition, we removed any regions that appeared unstructured. The remaining consensus sequence was then analyzed (**Figure 2**, blue regions). **Table S1** shows the aligned sequences of all structures used in this study. Regions of removed sequence as well as any other resolved proteins or peptides were implicitly included in our model by means of vibrational subsystem analysis (**VSA**) [38]. Any stabilizing antibody chains resolved in the structure were not included in the calculations.

We analyzed a total 26 X-ray structures. Of these, 9 were apo structures, 1 was clathrin-bound, 2 were inositol-6-phosphate (IP6) activated structures, 4 were GPCR-activated arrestin structures, and 10 were peptide-activated (**Table 1**) [39–51]. Nineteen (19) of the structures were arrestin-2, while 7 were arrestin-3. ANM was then performed using the alpha carbons of all 26 structures using VSA to model in all removed sequences, and the resulting eigenvalues and eigenvectors were saved. This resulted in our final data set.

**Table 1:** Arrestin Structures

| PDB ID | PMID | Arrestin | Type |
| --- | --- | --- | --- |
| 3P2D | 21215759 | 3 | Apo |
| 1G4R | 11566136 | 2 | Apo |
| 2WTR |  | 2 | Apo |
| 6KL7 | 31948726 | 2 | Apo |
| 1G4M | 11566136 | 2 | Apo |
| 1JSY | 11876640 | 2 | Apo |
| 3GD1 | 19710023 | 2 | Clathrin-Bound |
| 1ZSH | 16439357 | 2 | IP6 |
| 5TV1 | 29127291 | 3 | IP6 |
| 6UP7 | 31945771 | 2 | GPCR-bound |
| 6U1N | 31945772 | 2 | GPCR-bound |
| 6TKO | 32555462 | 2 | GPCR-bound |
| 6PWC | 31776446 | 2 | GPCR-bound |
| 6NI2 | 31740855 | 2 | Vasopressin-Bound |
| 4JQI | 23604254 | 2 | Vasopressin-Bound |
| 7DF9 | 33888704 | 2 | Vasopressin-Bound |
| 7DFA | 33888704 | 2 | Vasopressin-Bound |
| 7DFB | 33888704 | 2 | Vasopressin-Bound |
| 7DFC | 33888704 | 2 | Vasopressin-Bound |
| 6K3F | 32579945 | 3 | CXCR7-Bound |

### 3.2 Anisotropic Network Modeling

ANM models the protein as a network of beads connected by springs, with each bead representing the position of a Cα atom. The potential energy between the i^th^ and j^th^ Cα in the network is given by Hooke’s Law:

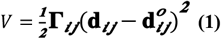

where 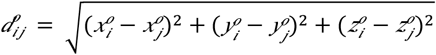 is the distance between the i^th^ and j^th^ Cα atom in the reference structure, and Γ_*ij*_ is the spring constant between them [28, 30, 52]. As a result, in this formalism the reference structure is the global minimum energy state. The spring constant is defined as:

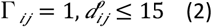

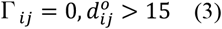

This creates a 3N x 3N Hessian matrix, where N is the number of nodes in the network (Cα atom in the structure). When diagonalized, the matrix returns eigenvalues (λ*i*) and eigenvectors 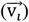 corresponding to the vibrational modes of the protein. The eigenvectors are the directions of motion, with each associated eigenvalue corresponding to the frequency of that motion. In ANM, the frequency of motion is inversely proportional to the amplitude of motion, due to the model’s harmonic nature. This means that the lowest frequency modes represent the highest amplitude motions. The 6 zero frequency modes corresponding to rigid body translation and rotation are ignored for all subsequent analysis.

The various structures used in our model had differennt numbers of Cα atoms, and the resulting eigendecompositions can only be easily compared when the matrix dimensions are identical. To accomplish this we used VSA [29, 31, 38] as implemented in LOOS [29, 53]. This method partitions the Hessian matrix into an environment and a subsystem, where the subsystem contains all consensus residues; these are the amino acids for which the vibrational motions are computed. The remaining residues are part of the environment, and their effects on the subsystem are included implicitly. This approach allows us to use the same subsystem for all proteins, which facilitates direct comparison, while still including the rest of the resolved structure.

### 3.2 Covariance complement

We compared our ANM eigenvectors and eigenvalues of the various arrestins to each other using a modified version of covariance overlap, called covariance complement [30, 31, 54].

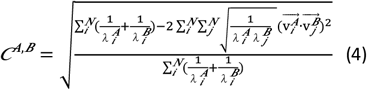

Where 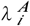 and 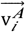 are the i^th^ eigenvalue and eigenvector of structure A, and 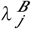 and 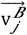 are the j^th^ eigenvalue and eigenvector of structure B. The covariance complement is 0 when the two ANM eigensets are identical and 1 when they are completely orthogonal. The primary advantage of using the covariance overlap or complement is that they compare the entire eigenset (directions of motion and amplitudes), rather than an arbitrarily chosen subset.

### 3.3 Per-Residue Contribution to Dot Product

We compared the low modes of our ANM eigenvectors to determine how much each residue contributes to the overall difference to the total dot product. We define this as a vector where each residue r consists of a 3-dimension sub-vector taken from the eigenvector used in equation 4. This represents the 3-dimensional motions of each individual motion of that residue as defined by the eigenvector. We then compute a difference score 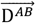 by:

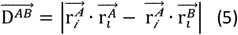

where 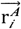 is the (x,y,z) components of a specific eigenvetor corresponding to the motions of the ith residue of structure A and 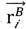 is the equivalent from structure B. Since the original eigenvector has already been normalized, the sum of the dot product of all residues, 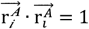, and residues 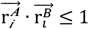. We then computed the average 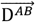 by averaging over all possible combinations of structures A and B and present the data in arbitrary units based on the contribution.

## 4. Results

### 4.1 Structures and Dynamics Comparison

There are several crystallographic structures of both arrestin-2 and arrestin-3. These structures include apo arrestin [40, 44, 47, 50, 51], as well as arrestin bound to clathrin [43], inositol hexaphosphate (**IP6**) [39, 46], activating peptides [41, 55, 56], and GPCRs [42, 48, 49]. They form three distinct structural clusters, as shown by Figure 3: an apo cluster with the C-tail domain bound to the N-domain, and two distinct active conformations representing the C-tail unbinding and the N-domain and C-domain rotating with respect to each other. Arrestin-2 and arrestin-3 form two distinct structures that differ mainly by the degree of rotation between the N-domain and C-domains [39].

**Figure 3:**
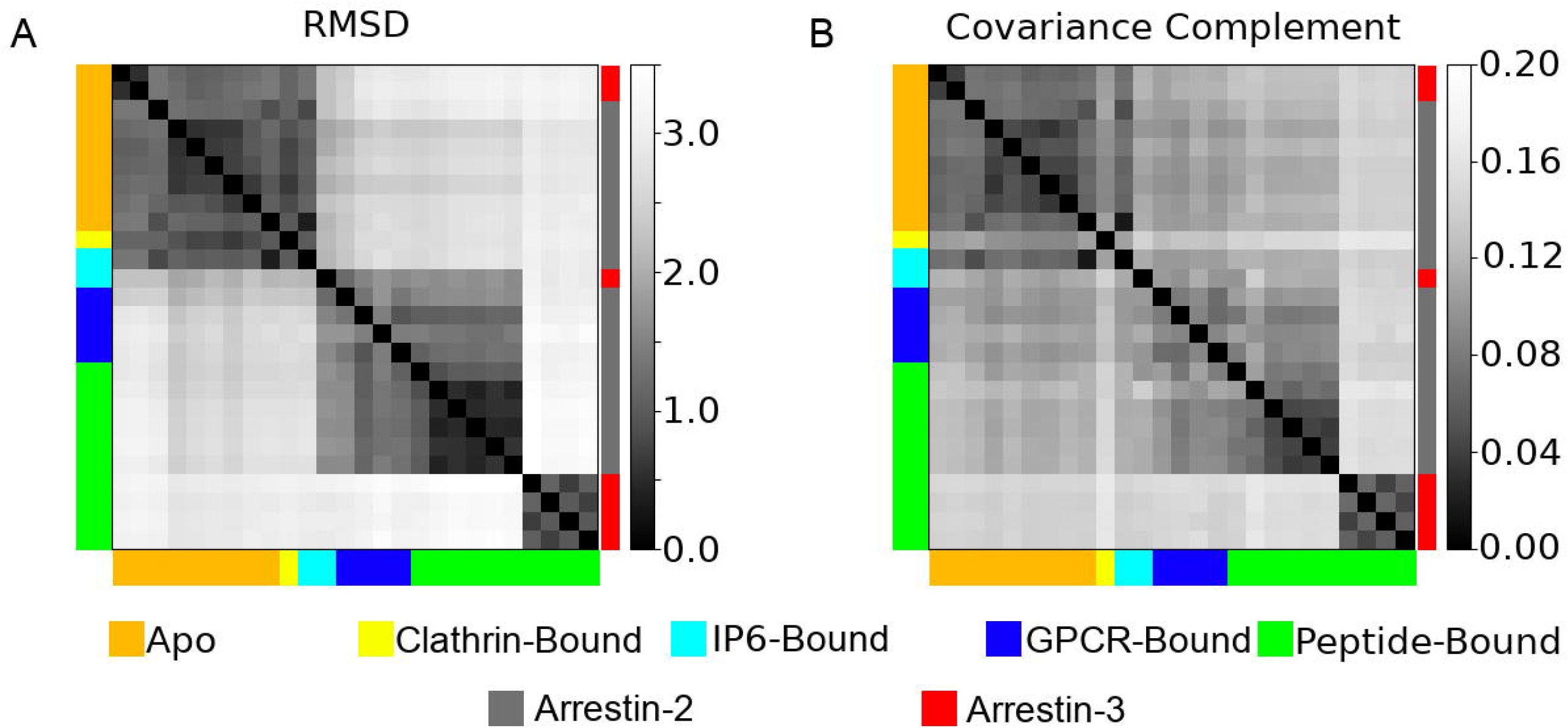
A pairwise comparison of the A) root mean squared deviation (RMSD) and B) the covariance complement derived from the 26 different structures analyzed in this study. Structures are color coded both by their ligands (left and bottom of graph), or whether they are arrestin-2 (gray) or arrestin-3 (red) (right of graph).

We analyzed 26 protein chains from 20 distinct crystal structures (**Table 1**); Figure 2 shows their common structural elements. We compared their structure using the standard alpha-carbon RMSD after alignment. We used the covariance complement (**Eq 4**) to quantify the differences in their fluctuations, as estimated using VSA; the results are shown as Figure 3.

The first take-home message is that all structures are quite similar, with a maximum pairwise RMSD of 3.46 and covariance complement of 0.149. That said, the structures form some distinct clusters readily visible in the heatmaps. For example, all apo arrestin-2 and arrestin-3 structures are extremely similar; this cluster also contains the clathrin-bound structure, 3GD1, and one of the IP6-bound structures, 1ZSH. The GPCR-bound and peptide-bound arrestin structures form 2 distinct clusters. The first contains the arrestin-2 structures and the other IP6-bound structure, 5TV1, while the second contains the arrestin-3 structures (**Figure 3A**). The main difference between these clusters is the degree of rotation between the N-domain and the C-domain.

Focusing on the protein fluctuations (**Figure 3B**) tells a subtly different story. The apo structures and peptide-bound arrestin-3 fluctuations form two clusters corresponding to those seen in the structural analysis. However, the single clathrin-bound structure has dynamics that place it essentially in a cluster by itself, despite its structural similarity to the apo proteins. The vasopressin-bound arrestin-2 structures (6U1N, 6NI2, 4JQI, 7DF9, 7DFA, 7DFB, 7DFC), which structurally clustered with the other peptide-bound and GPCR-bound chains, form their own distinct cluster from a dynamics perspective. The remaining arrestin-2 chains and the other IP6-bound chain appear to be as dissimilar from each other as they are from the apo structures (**Figure 3B**). From this, we conclude activated arrestins have similar structures but unique dynamics that are determined by the peptide they are bound to.

### 4.2 Comparison of Modes of Motion

We did a mode-wise comparison of the various eigensets and discovered that the first two modes of all eigensets were very similar, as shown in Figure 4; mode 3 also showed some similarity (**Figure S1**), but beyond that mode-mixing meant similarity was low. For mode 1, with the exception of the clathrin-bound structure (3GD1), the absolute dot products were mostly above 0.9, indicating that the modes are nearly parallel. The set of first modes naturally form 2 clusters; the arrestin-3 structures cluster with the apo and IP6-bound ones, in contrast to the simple RMSD measurement, which puts arrestin-3 in its own cluster. The second cluster contains the peptide-bound and GPCR-bound arrestin-2 structures. However, despite the existence of visually identifiable clusters, all of the mode 1’s are strikingly similar; the farthest outlier (3GD1) still produces a mode 1 with an average absolute dot product of 0.68 with the other structures. While far lower than the others, this value would be virtually impossible to get by chance with random vectors in this dimension (**Figure S2**). Still, the results suggest that clathrin binding alters the dynamics of the protein. Mode 2 (**Figure 4b**) tells a virtually identical story, although the similarity between the two clusters (between the peptide-and GPCR-bound arrestin-2s and the rest) is a little lower.

**Figure 4:**
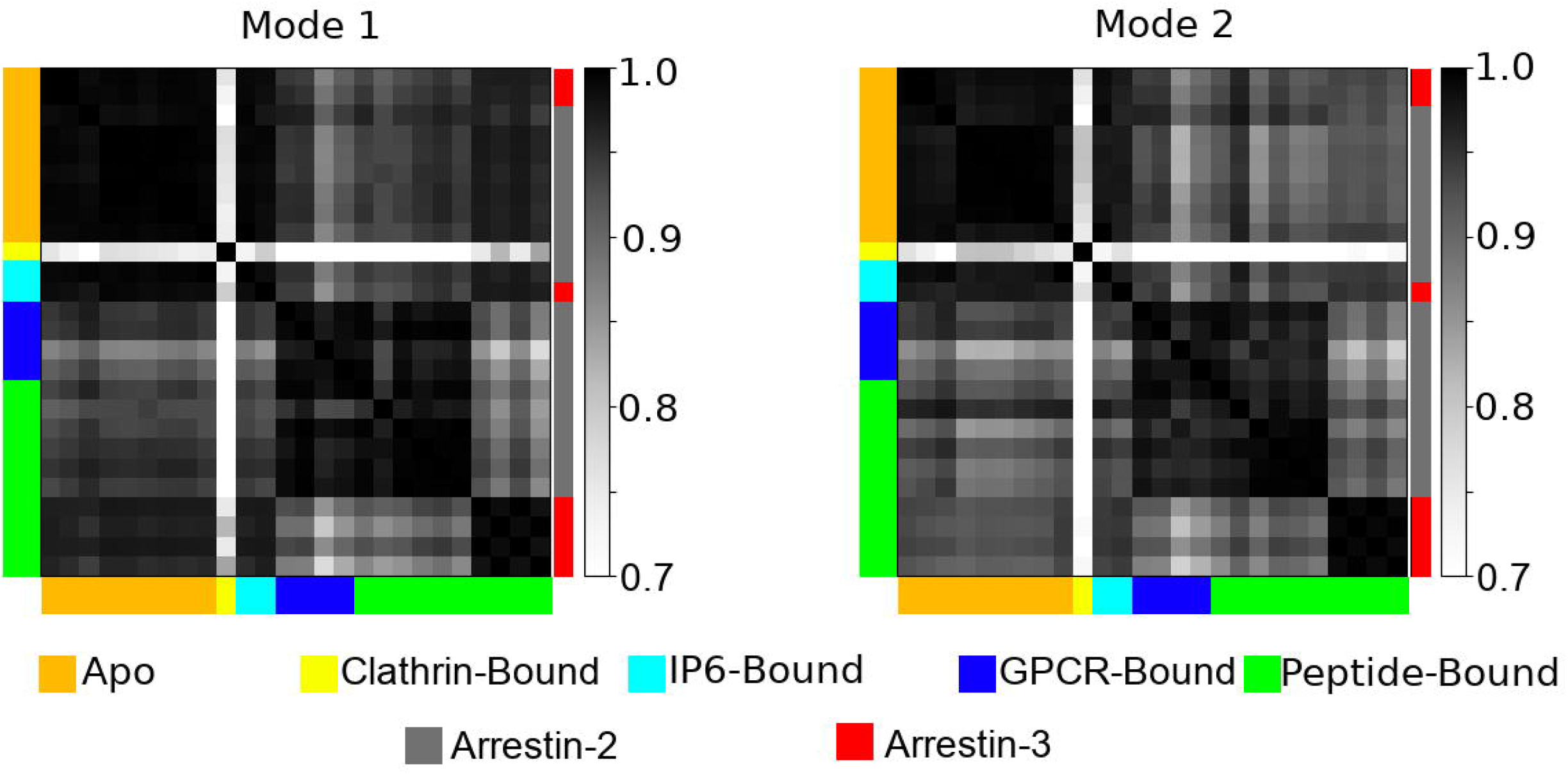
A pairwise dot product of the first and second modes of motion as computed by ANM for all 26 structures analyzed. Structures are color coded both by their ligands (left and bottom of graph), or whether they are arrestin-2 (gray) or arrestin-3 (red) (right of graph).

### 4.3 Comparison of Activated to Apo Arrestins

Although overall quite similar, the lowest modes for apo arrestins cluster separately from the activated arrestin-2 structures (**Figure 4**). This suggests that there are common changes to dynamics upon arrestin-2 activation by either a phosphopeptide or a GPCR but not IP6. We visualized this change in mode one by taking example structures from the GPCR-bound (6TKO) and apo (2WTR) arrestin-2 and mapping their respective mode 1s onto the structure of apo arrestin (**Figure 5**). This reveals that most of the difference comes from a change in the motion of α-helix 1 and the C-domain β-sheets on the C-loop side.

**Figure 5:**
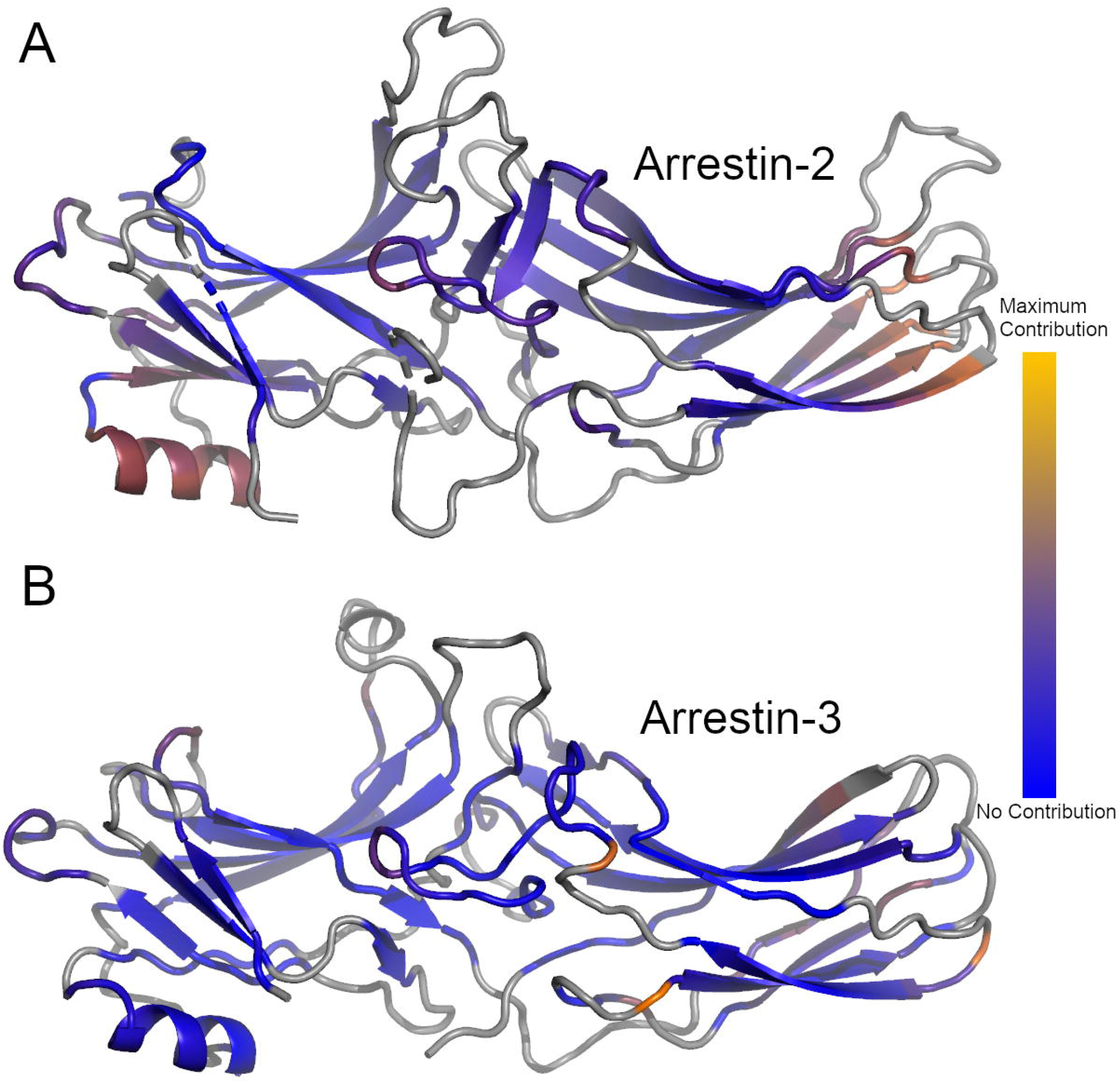
Average difference in the per-residue dot product between the first mode of all unliganded and peptide-bound structures for Arrestin-2 (A) or Arrestin-3 (B). Gray regions represent structural elements not included in the ANM model. Blue regions represent regions which did not contribute to the differences in the dot product, orange regions represent regions of maximal contribution.

This difference can be quantified by computing the average amplitude of the difference between the respective first modes on a per-residue basis. Figure 5a shows the results by color-coding the structure of arrestin-2 using this quantity, which naturally highlights α-helix 1 and the C-domain β-sheets. Interestingly, this pattern is not retained when peptide-activated arrestin-3 is compared to the apo structures. Figure 5b shows that helix 1’s motions are similar in the two sets, and while there are differences in the C-domain β-sheet, they are smaller and involve fewer residues. We can further visualize the changes in these changes by looking at the change in the vector of motion between two typical structures (2WTR/6TKO) and visualizing the change in their first mode of motion (**Figure S3**).

The other area where mode 1 differs between apo and activated arrestin-2 and arrestin-3 is the middle portion of the lariat loop, shown in the center of both structures. In the arrestin-3 structure shown, these residues are in contact with the bound peptide, making a dynamic change less surprising; the origin of the change in arrestin-2 is less clear.

### 4.4 Comparison of Apo to Clathrin-bound Arrestins

The lone clathrin-bound structure is structurally nearly identical to apo arrestin-2 and arrestin-3 with an average RMSD of 0.66 Å from all apo arrestin structures. Despite this, it shows the most dramatic differences in its low mode dynamics compared to all other structures (**Figure 4**). This suggests that binding partners to arrestin can dramatically affect its dynamics without visibly perturbing its structure.

We mapped the average per-residue difference between the amplitudes of the first mode of all apo structures to clathrin bound arrestin-2 and we plotted these changes onto the structure of clathrin-bound arrestin-2 (3GD1) (**Figure 6**). As with comparing apo and activated arrestin-2 the differences in the motion are concentrated in α-helix 1 and the C-loop side of the C-domain β-sheets. The effects on the lariat loop are minimal, and there is an additional perturbation in motions in the semi-structures N-domain loop 44D-52R that does not appear in the differences between apo and active arrestin.

**Figure 6:**
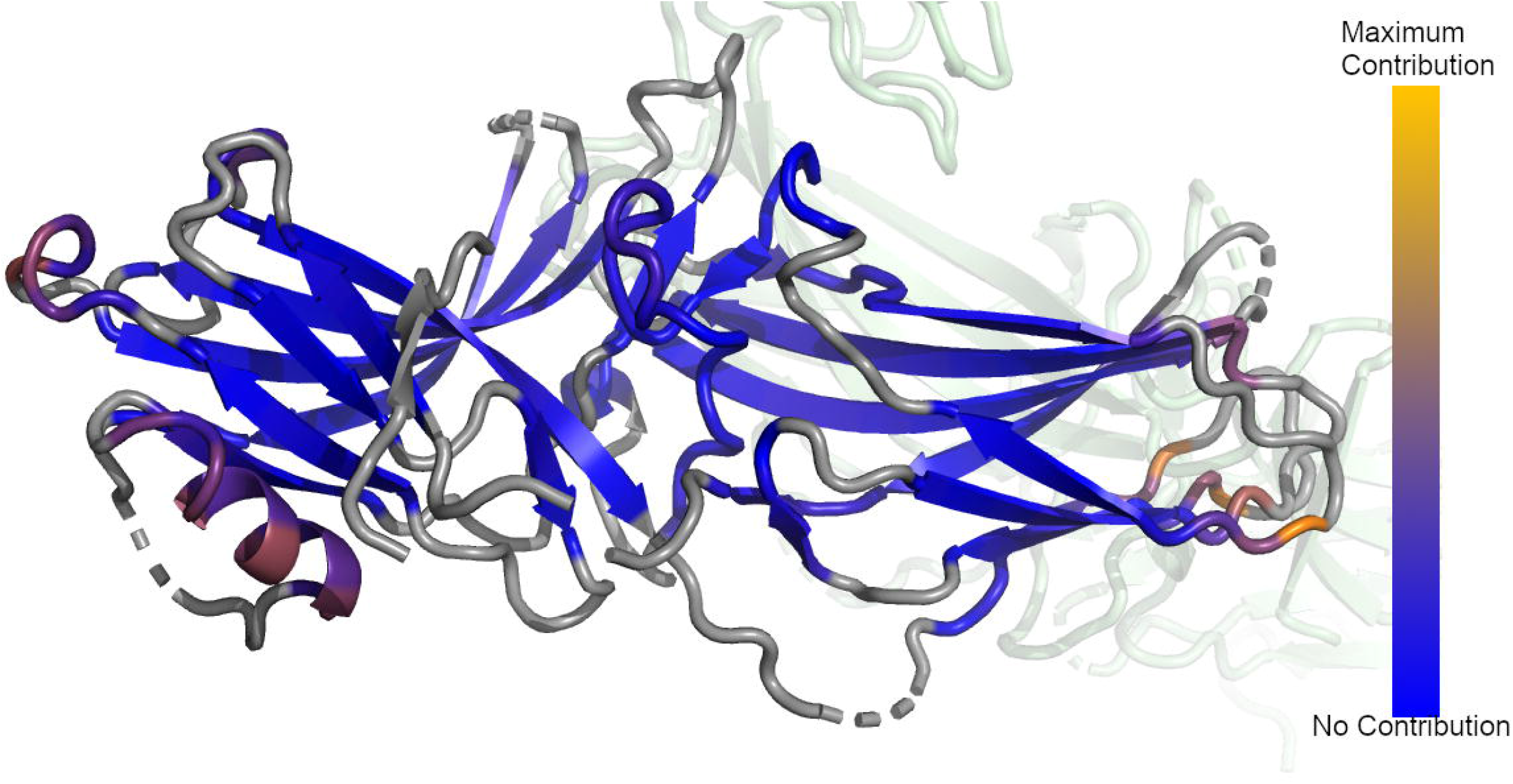
Average difference in the per-residue dot product between the first mode of all unliganded and clathrin-bound Arrestin-2. Gray regions represent structural elements not included in the ANM model. Blue regions represent regions which did not contribute to the differences in the dot product, orange regions represent regions of maximal contribution.

We also examined the difference between the second mode of apo and clathrin bound arrestin-2 and found the differences were localized to the same regions (data not shown). Furthermore, inspection of the difference between active arrestin and clathrin bound arrestin revealed the same regions changing their motions.

## 5. Discussion

### 5.1 Comparison To Experimental Studies

It is well known that arrestin’s structure and dynamics change upon activation [26, 57]. However, the precise nature of these changes, particularly in dynamics, and their relation to activity remains elusive. This has been the focus of numerous crystallographic studies [39–44, 46–51]. There also have been several HX-MS and electron paramagnetic resonance (**EPR**) studies that probe the solution structure and dynamics of arrestin and how those dynamics change upon activation [6, 22, 44, 57–62]. The aim of the present study is to use the existing crystallographic data to predict how structural changes upon activation change the resulting dynamics. Our work provides insights into two different stages of arrestin activation: (1) How arrestin dynamics change upon activation by a phosphopeptide or GPCR, and (2) how arrestin dynamics vary with subtle changes to the activating peptides phosphorylation status.

It is well established that the lowest modes produced by network models correlate well with HX-MS results [33, 35, 63]. There have been multiple HX-MS studies that probed the structural dynamics of apo wild-type arrestin, as well mutants that allow arrestin to assume an active-like conformation in the absence of a GPCR or activating peptide [6, 57, 59, 61], where these pre-activated mutants generally disrupt the charge balance within the polar core [64–66].

### 5.2 Low mode differences between active and apo arrestin

Our model predicts that arrestin-2’s motions change significantly upon activation by peptide or GPCR binding (**Figure 5**). We found that α-helix 1, the β-sheets bordering the C-edge loops, and L293 and 294K of the lariat loop of arrestin-2 show the greatest change in their dynamics as measured by the lowest mode of the ANM eigensets. These regions are of particular interest since they are three functionally important regions for arrestin. Vishnivetskiy et. al. showed that α-helix 1 plays a role in receptor binding, and that mutations to α-helix 1 disrupt receptor binding in arrestin-1 [22]. It is not surprising that this region shows up in our analysis as well, since this is precisely where the ligand (not present in the apo structures) binds. Meanwhile, the C-edge loops have been shown to bind to both clathrin and the cell membrane upon activation, and our observed change in dynamics may play a role in this [9, 10, 27]. Lysine 294 is in the middle of the lariat loop and binds phosphorylated residues on the GPCR; its homolog in arrestin-3 contacts bound IP6 [39, 41]. However, while it plays a role in receptor binding it does not serve as a phosphosensor [67]. As a result, we believe the calculations identify a set of allosterically linked sites on arrestin-2 that change their motions to enhance receptor binding and aid in endocytosis by simultaneously enabling membrane binding by coupling the motions of the GPCR activating peptide binding site with functional domains such as the C-loops which aid in membrane binding [27].

Activated arrestin-3 shows significantly less difference in its lowest mode dynamics compared to apo arrestin-3. Most of the residues identified appear to be isolated changes, although there are a cluster of dynamics changes in the β-sheets bordering the C-edge loops, and residues L295 and K296 of the lariat loop. However, α-helix 1 is conspicuously absent from this change. This suggests that arrestin-3 does not rely on α-helix 1 to enhance receptor binding, although its modest changes to dynamics upon activation may enhance membrane binding; it is worth noting that any interactions with the membrane are not included in the ANM model.

### 5.3 Low mode differences between clathrin-bound arrestin-2 and apo arrestin

Clathrin-bound arrestin-2 shows a similar fingerprint of differences when compared to both apo and active arrestin-2 with the exception of N-domain loop 44D-52R, which contains the NLS for vasopressin-bound arrestin-2 (P45-R51).[41] This region has also been experimentally shown to change its structure based on the phosphorylation pattern of activating peptide bound to arrestin-2.[41] This fits well with the idea that clathrin binding to arrestin triggers internalization and suggests that this occurs at least in part by means of an allosteric interaction between the clathrin binding site and the NLS.[9]

### 5.4 Identification of an allosteric network in arrestin

Our data reveals several disparate regions which include α-helix 1, the β-sheets bordering the C-edge loops, the lariat loop, and the NLS which appear to change both the amplitude and direction of their normal modes of motion in response to any sort of perturbation. These sites change their motions in response to clathrin binding, even though this binding does not disrupt arrestin’s polar core and hence does not make it assume its active conformation. The fact that these sites are allosterically linked could give a clue as to how arrestin works within the cell.

### 5.5 Conclusion

We have performed a survey of all available structures of arrestin-2 and arrestrin-3 using ANM. We have shown that all apo structures of arrestin-2 and arrestin-3 have nearly identical dynamics, but their dynamics change dramatically upon activation. Arrestin-2 and arrestin-3 assume markedly different dynamics from each other upon activation, but these dynamics are nearly as dissimilar from each other as they are from apo arrestin. Examination of the lowest modes of motion in these models reveals that the differences are confined to a small set of functionally important regions in arrestin-2 but not arrestin-3. This change in dynamics upon activation could explain some of arrestin-2 activity binding active GPCRs.

## Acknowledgements

This work was supported by NIH grant U01DA051373.

## Statements and Declarations

The authors have no financial or non-financial interests to declare

## Notes

<u>Data Availability Statement</u> The data that support the findings of this study are openly available at https://github.com/GrossfieldLab/arrestin-anm.

### Competing Interest Statement

The authors have declared no competing interest.

