## Supplementary figures and images for "Surveying non-visual arrestins reveals allosteric interactions between functional sites"

### Figure S1

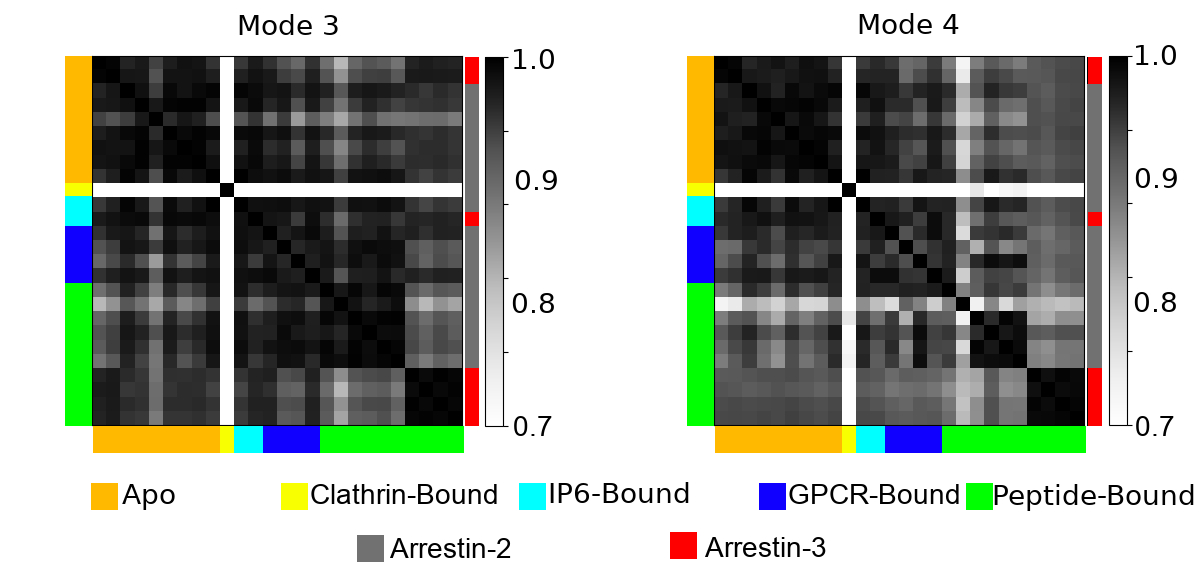

### Figure S2

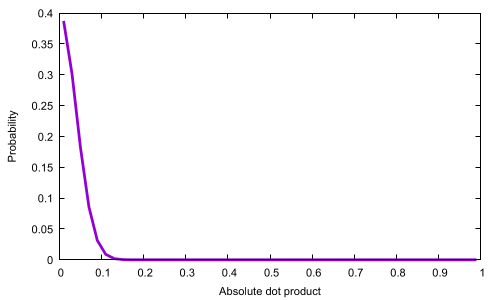

### Figure S3

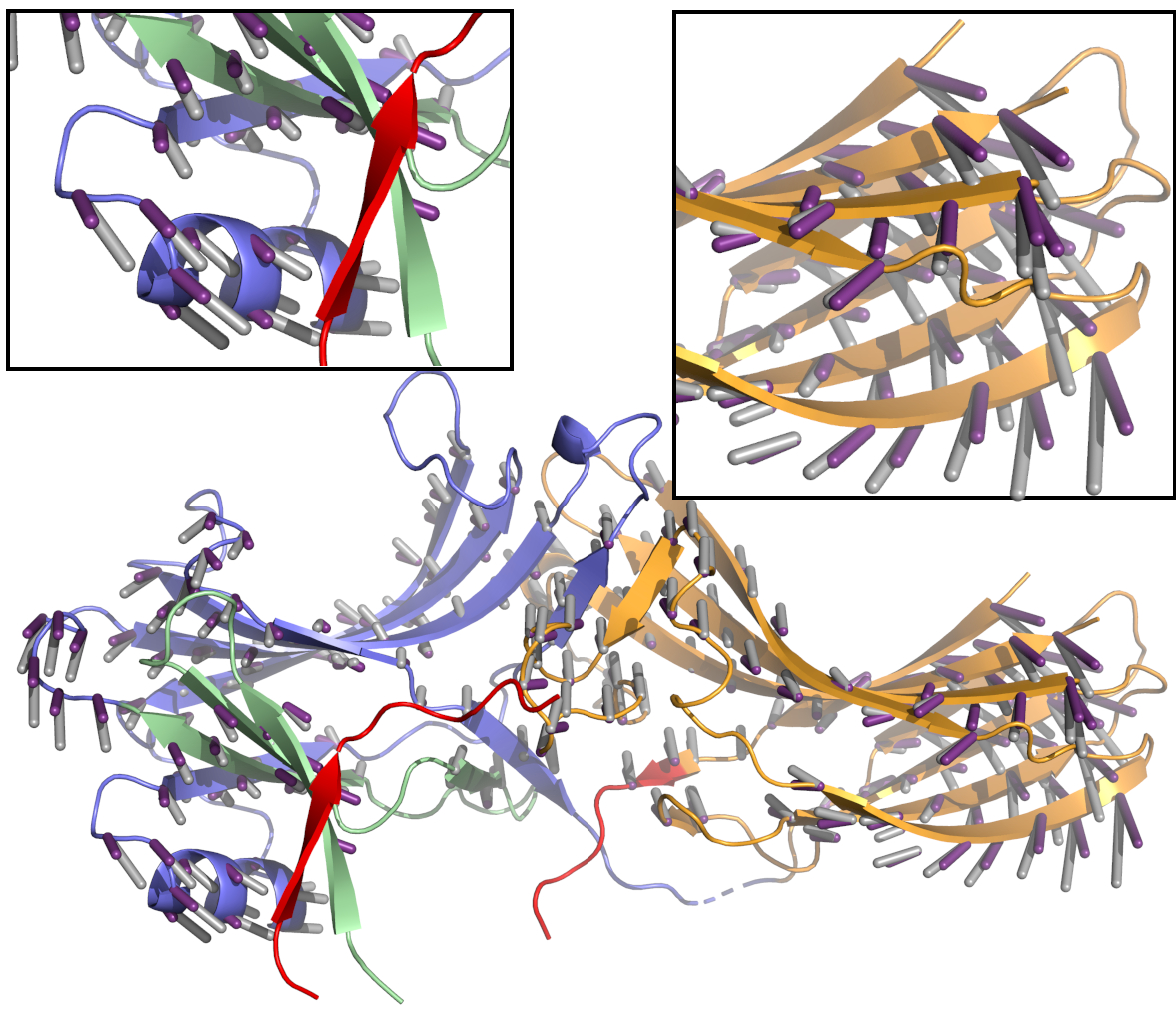
